# Most Compositae (Asteraceae) are descendants of a paleohexaploid and all share a paleotetraploid ancestor with the Calyceraceae

**DOI:** 10.1101/043455

**Authors:** Michael S. Barker, Zheng Li, Thomas I. Kidder, Chris R. Reardon, Zhao Lai, Luiz O. Oliveira, Moira Scascitelli, Loren H. Rieseberg

**Affiliations:** Department of Ecology & Evolutionary Biology, University of Arizona, Tucson, AZ 85721 USA; Department of Biology and Center for Genomics and Bioinformatics, Indiana University, Bloomington, IN 47405 USA; Departamento de Bioquímica e Biologia Molecular, Universidade Federal de Viçosa, Viçosa, Brazil; Department of Botany and Biodiversity Research Centre, University of British Columbia, Vancouver, BC, Canada

## Abstract

**Premise of the study:** Like many other flowering plants, members of the Compositae (Asteraceae) have a polyploid ancestry. Previous analyses found evidence for an ancient duplication or possibly triplication in the early evolutionary history of the family. We sought to better place this paleopolyploidy in the phylogeny and assess its nature.

**Methods:** We sequenced new transcriptomes for *Barnadesia*, the lineage sister to all other Compositae, and four representatives of closely related families. Using a recently developed algorithm, MAPS, we analyzed nuclear gene family phylogenies for evidence of paleopolyploidy.

**Key results:** We found that the previously recognized Compositae paleopolyploidy is also in the ancestry of the Calyceraceae. Our phylogenomic analyses uncovered evidence for a successive second round of genome duplication among all sampled Compositae except *Barnadesia*.

**Conclusions:** Our analyses of new samples with new tools provide a revised view of paleopolyploidy in the Compositae. Together with results from a high density *Lactuca* linkage map, our results suggest that the Compositae and Calyceraceae have a common paleotetraploid ancestor and most Compositae are descendants of a paleohexaploid. Although paleohexaploids have been previously identified, this is the first example where the paleotetraploid and paleohexaploid lineages have survived over tens of millions of years. The complex polyploidy in the ancestry of the Compositae and Calyceraceae represents a unique opportunity to study the long-term evolutionary fates and consequences of different ploidal levels.

## INTRODUCTION

Polyploidy, or speciation by whole genome duplication (WGD), is widespread throughout the history of vascular plants. Nearly 15% of flowering plant speciation events are caused by polyploidy (Wood et al., 2009) and approximately 30% of plant species are recent polyploids (Wood et al., 2009; Mayrose et al., 2011; Barker et al., 2016). Numerous paleopolyploidies have been described among and within families (Schlueter et al., 2004; Cui et al., 2006; Jaillon et al., 2007; Barker et al., 2008, 2009; Rensing et al., 2008; Tang et al., 2008; Soltis et al., 2009; Schmutz et al., 2010; Shi et al., 2010; Velasco et al., 2010; Wang et al., 2011; Arrigo and Barker, 2012; Jiao et al., 2012, 2014; Tomato Genome Consortium, 2012; Vekemans et al., 2012; Cheng et al., 2013; Ibarra-Laclette et al., 2013; Ming et al., 2013; Estep et al., 2014; Sveinsson et al., 2014; Cai et al., 2015; Cannon et al., 2015; Edger et al., 2015; Li et al., 2015; Marques et al., 2016; Yang et al., 2015), as well as in the ancestry of all seed plants (Jiao et al., 2011; Li et al., 2015). Although we know the approximate phylogenetic placement of many paleopolyploidies, we often understand less about their nature. The formation of polyploid species may involve duplication of the genome(s) from one or more species. The outcomes include autopolyploids, allopolyploids, and a series of intermediates such as segmental allopolyploids (Kihara and Ono, 1926; deWet, 1980; Ramsey and Schemske, 1998; Barker et al., 2016). Crosses among taxa of different ploidal levels may lead to hexaploids and other higher level polyploids. Given the potential for polyploids of various natures to have different evolutionary outcomes (Otto and Whitton, 2000; Doyle et al., 2008; Hegarty and Hiscock, 2008; Soltis et al., 2010; Arrigo and Barker,2012; Buggs et al., 2014; Crawford et al., 2014; Garsmeur et al., 2014; Barker et al., 2016), a better characterization of paleopolyploidies is an important next step in understanding the evolution of plant genomes.

Analyses of well assembled plant nuclear genomes have revealed the nature of some ancient polyploidies. Ploidal level has been inferred by searching for evidence of multiplied blocks of collinear genes in syntenic data from whole genome sequences or linkage mapping studies. For example, paleohexaploidy has been inferred in the ancestry of the core eudicots by identifying triplicated blocks of collinear genes (Tang et al., 2008). Similar evidence for paleohexaploidy has been inferred in the Solanaceae (Tomato Genome Consortium, 2012) and the tribe Brassiceae (Lagercrantz and Lydiate, 1996; Lysak et al., 2005; Tang et al.,

2012). A recent high density linkage map of *Lactuca sativa* L. found evidence that the polyploidy in the ancestry of Compositae (Asteraceae) (Barker et al., 2008) may be a paleohexaploidy (Truco et al., 2013). Truco et al. (2013) observed that the *L. sativa* genome was triplicated when compared to the *Vitis vinifera* L. genome. However, their analyses could not phylogenetically place the set of genome duplications that generated the *Lactuca* paleohexaploidy. Using new data and phylogenomic methods, we more precisely placed this complex polyploidy in the history of the Compositae and closely related families.

Although most previous methods for identifying paleohexaploidy have used some form of syntenic or linkage data, a gene family approach with sufficient taxon sampling may also disentangle ancient hexaploidy and tetraploidy. Instead of identifying triplicated blocks of collinear genes, successive rounds of WGD should appear as phylogenetically nested duplications in nuclear gene family phylogenies. Higher ploidal level polyploidies, such as hexaploids, are formed by some combination of lower ploidal levels with genome doubling (Harlan and DeWet, 1975; Ramsey and Schemske, 1998). Hexaploids may be formed when a tetraploid crosses with a diploid. Triploids produced by these crosses may produce hexaploids when crossed themselves via unreduced 3*x* gametes. Alternatively, hexaploids may be formed from crosses between reduced (2*x*) and unreduced (4*x*) gametes from tetraploids with no diploids involved. Perhaps more rare are crosses of unreduced diploid (2*x*) and unreduced tetraploid (4*x*) gametes that may give rise to hexaploids. All of these pathways to hexaploids involve two, successive rounds of WGD that we may detect in the phylogenies of gene families from multiple species. Studying polyploidies that occurred tens of millions of years ago, it may be difficult to identify successive rounds of duplication if the WGDs happened very close in time or if the involved tetraploid or diploid lineages went extinct. However, if we have sampled extant representatives of the diploids or tetraploids involved in hexaploid formation, then we may place and infer the consecutive rounds of polyploidy giving rise to a paleohexaploid genome.

Using new data and algorithms, we re-evaluated our previous placement of the paleopolyploidy putatively shared by all extant Compositae (Barker et al., 2008). We sequenced new transcriptomes from the Barnadesieae (sister to all other Compositae), Calyceracae, Goodeniaceae, Menyanthaceae, and Campanulaceae. WGDs were inferred and placed in phylogenetic context using single species Ks plots of gene duplications (Barker et al., 2008, 2010) and our recently developed MAPS algorithm (Li et al., 2015). We also used MAPS to evaluate evidence of nested WGDs among our sampled taxa consistent with a paleotetraploidy followed by a paleohexaploidization.

## MATERIALS AND METHODS

### Transcriptome sampling–

We sequenced and assembled new transcriptomes for five species of Compositae and related families. Transcriptomes were sequenced for *Barnadesia spinosa* L.f. (Compositae), *Acicarpha spathulata* R.Br. (Calyceraceae), *Scaevola aemula* R.Br. (Goodeniaceae), *Nymphoides peltata* (S.G. Gmel.) Kuntze (Menyanthaceae), and *Platycodon grandiflorus* A.DC. (Campanulaceae). *Barnadesia* was sequenced using Sanger technology, whereas the other transcriptomes were sequenced using 454 technology. RNA was extracted from leaves, flower buds, and mature flowers from each species. We isolated RNA using either Trizol reagent (Invitrogen, Carlsbad, CA), RNeasy kits (Qiagen, Valencia, CA), or a combination of the two methods. In the combined approach, the standard Trizol protocol was followed through the chloroform extraction step at which point 0.53X volumes of 100% ethanol was added to the aqueous phase and the entire RNA/ethanol mixture then applied to an RNeasy column and the Qiagen protocol followed thereafter. Approximately equal amounts of total RNA isolated from each tissue type were pooled prior to EST library preparation.

Two different methods were used to generate EST libraries. For Sanger sequencing, we prepared normalized libraries with the TRIMMER-DIRECT cDNA Normalization Kit (Evrogen, Moscow, Russia). cDNA samples from both the standard and normalized EST libraries were size-fractionated through agarose gels into three classes (0.5 - 1 kb, 1 – 2 kb, 2 – 3 kb) to reduce size biases during the subsequent cloning and sequencing steps. For 454 sequencing (454 Life Sciences, Branford, CT), we used modified oligo-dT primers during cDNA synthesis to reduce the length of mononucleotide runs associated with the polyA tail of mRNA. Mononucleotide runs reduce sequence quality and quantity due to excessive light production and crosstalk between neighboring cells. For cDNA synthesis of the 454 libraries, we either used a broken chain short oligo-dT primer (Meyer et al., 2009) or two different modified oligo-dT primers: one to prime the polyA tail of mRNA during first strand cDNA synthesis and another to further break down the stretches of polyA sequence during second strand cDNA synthesis (Beldade et al., 2006; Lai, Zou, et al., 2012). We then normalized and amplified the cDNA using the TRIMMER-DIRECT cDNA Normalization Kit as above. After normalization, cDNA was sonicated to 500 to 800bp fragments and size-selected to remove small fragments. Then the fragmented ends were polished and ligated with adaptors. The optimal ligation products were selectively amplified and subjected to two rounds of size selection including gel electrophoresis and AMpure SPRI bead purification.

The *Barnadesia* Sanger EST library was sequenced using ABI 3730 machines (Life Technologies, Carlsbad, CA) at the Joint Genome Institute, Walnut Creek, CA. All other 454 libraries were sequenced on GS-FLX machines at Genome Quebec (http://www.genomequebec.com) using the standard Titanium Chemistry (http://454.com/). Raw reads for each newly sequenced transcriptome were cleaned and assembled into contigs. We used the SnoWhite cleaning pipeline (Dlugosch et al., 2013) to remove adapters, chimeras, and other possible contaminating sequences from raw reads. The cleaned reads were assembled using MIRA with the recommended settings for Sanger or 454 assemblies (Chevreux et al., 2004). Following MIRA, we finished the transcriptome assemblies with CAP3 (Huang and Madan, 1999) using a percent overlap of 94%. We also downloaded Sanger sequenced EST data from GenBank for *Artemisia annua* L. (Nair et al., 2013) and assembled with the same pipeline. Our previous transcriptome assemblies for *Helianthus annuus* L., *Lactuca sativa* L., and *Centaurea solstitialis* L. (Barker et al., 2008; Lai, Kane, et al., 2012) were also used in this analysis. All raw and assembled data are available through the Compositae Genome Project Database(http://compgenomics.ucdavis.edu/compositae_database.php).

### DupPipe analyses of WGDs from transcriptomes of single species—

For each transcriptome, we used our DupPipe pipeline to construct gene families and estimate the age of gene duplications (Barker et al., 2008, 2010). We translated DNA sequences and identified reading frames by comparing the Genewise alignment to the best hit protein from a collection of proteins from 25 plant genomes from Phytozome (Goodstein et al., 2012). For all DupPipe runs, we used protein-guided DNA alignments to align our nucleic acids while maintaining reading frame. We estimated synonymous divergence (Ks) using PAML with the F3X4 model (Yang, 2007) for each node in our gene family phylogenies. We identified peaks of gene duplication as evidence of ancient WGDs in histograms of the age distribution of gene duplications (Ks plots). We used a mixture model, EMMIX (McLachlan et al., 1999). to identify significant peaks and estimate their median Ks values.

### MAPS analyses of WGDs from transcriptomes of multiple species—

To further infer and locate paleopolyploidy in our data sets, we used our recently developed gene tree sorting and counting algorithm, the Multi-tAxon Paleopolyploidy Search (MAPS) tool (Li et al., 2015)https://bitbucket.org/barkerlab/maps: (Li et al., 2015). The MAPS algorithm uses a given species tree to filter collections of nuclear gene trees for subtrees consistent with relationships at each node in the species tree. Using this filtered set of subtrees, MAPS identifies and records nodes with a gene duplication shared by descendant taxa. To infer and locate a potential whole genome duplication, we plotted the percentage of gene duplications shared by descendant taxa by node. A WGD will produce a large burst of shared duplications across taxa and gene trees. This burst of duplication will appear as an increase in the percentage of shared gene duplications in our MAPS analyses (Li et al.,2015). If the inferred WGDs are not too saturated, they will also appear in Ks plots for each species and corroborate the phylogenetic placement of WGDs. For our MAPS analysis, we used transcriptomes from five Compositae species and four outgroup species each representing closely related families of the Compositae. Our species tree for the MAPS analysis was based on previously published phylogenies (Kim et al., 2005; Panero and Funk, 2008; Funk et al., 2009; Tank and Donoghue, 2010; Soltis et al., 2011; Panero et al., 2014).

We circumscribed and constructed nuclear gene family phylogenies from multiple species for the MAPS analysis. We translated each transcriptome into amino acid sequences using the TransPipe pipeline as in our DupPipe analyses (Barker et al., 2010). Using these translations, we performed reciprocal protein BLAST (blastp) searches among data sets for the MAPS analysis using an E-value of 10e-5 as a cutoff. We clustered gene families from these BLAST results using OrthoFinder under the default parameters (Emms and Kelly, 2015). Using a custom perl script, we filtered for gene families that contained at least one gene copy from each taxon and discarded the remaining OrthoFinder clusters. We used PASTA for automatic alignment and phylogeny reconstruction of gene families (Mirarab et al., 2014). For each gene family phylogeny, we ran PASTA until we reached three iterations without an improvement in likelihood score using a centroid breaking strategy. We constructed alignments using MAFFT (Katoh et al., 2002), employed Muscle for mergers (Edgar, 2004), and RAxML for tree estimation (Stamatakis, 2014). The parameters for each software package were the default options for PASTA. We used the best scoring PASTA tree for each multi-species nuclear gene family to infer and locate WGDs using MAPS.

## RESULTS

Our newly sequenced and assembled transcriptomes for the Compositae and closely related families (Table 1) revealed a history of polyploidy. Peaks consistent with paleopolyploidy were observed in all Compositae and Calyceraceae species (Fig. 1). We previously identified WGD peaks ranging from Ks = 0.5–0.95 in the age distributions of *Centaurea, Helianthus,* and *Lactuca,* as well as an additional more recent WGD in the Heliantheae (Barker et al., 2008). The current analyses support these results and also find evidence for a WGD in the gene age distributions of *Artemisia* (Fig. 1A; median peak Ks = 0.86) and *Barnadesia* (Fig. 1B; median peak Ks = 0.56). Outside of the Compositae, we observed a similarly divergent WGD peak in the age distribution for *Acicarpha* from the Calyceraceae (Fig. 1C; median peak Ks = 0.56). The transcriptomes of the other outgroup taxa representing Goodeniaceae, Menyanthaceae, and Campanulaceae did not contain evidence of a WGD within a Ks < 2 window.

**Table 1.**
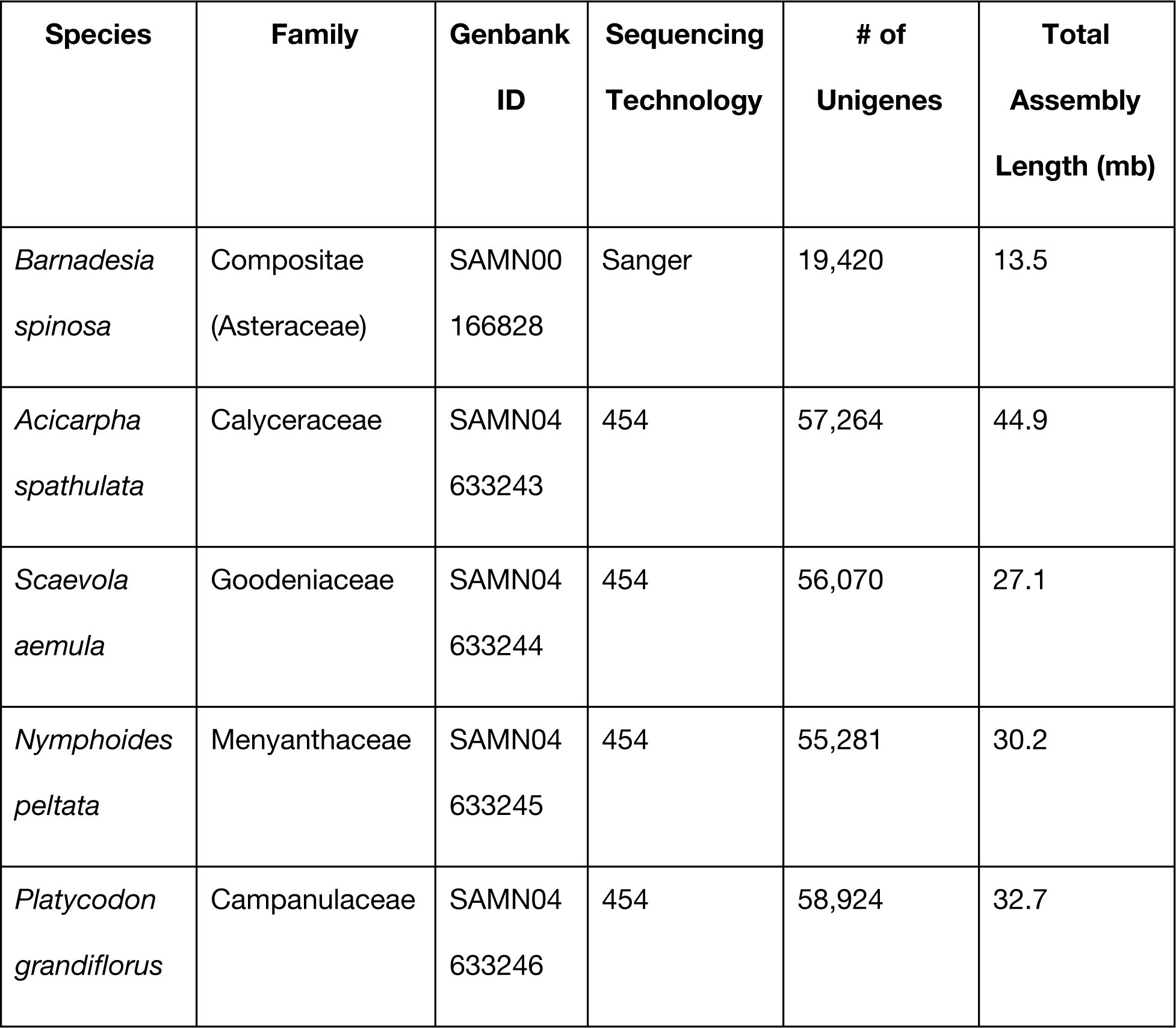
Summary statistics for the newly sequenced and assembled transcriptomes from species of Compositae and closely related families. All data available from the Compositae Genome Project Database (http://compgenomics.ucdavis.edu).

We used MAPS (Li et al., 2015) to test if the Compositae and Calyceraceae share an ancient WGD or experienced independent WGDs. Gene family clustering across the nine transcriptomes from species of Compositae and related families recovered 1,814 nuclear gene families with at least one copy from each sampled species. Among these gene families, MAPS revealed two bursts of shared gene duplication across the phylogeny (Fig. 2). One of these peaks of shared duplications was located in the MRCA of the Compositae and Calyceraceae. Forty percent of the gene subtrees consistent with the species tree supported a gene duplication event at this node (Fig. 2 & 3). A burst of shared gene duplications at this node is consistent with the observed pattern of peaks from the single species Ks plots; inferred WGDs in *Acicarpha, Barnadesia*, and all other Compositae, but no apparent WGDs in the representatives of other families (Fig. 1).

We also observed a second burst of shared gene duplications among the core members of the Compositae (Fig. 2 & 3). More than 45% of the gene subtrees supported a shared duplication in the MRCA of *Centaurea, Lactuca, Artemesia,* and *Helianthus.* Although not easily distinguished in the Ks plots, this second round of WGD is consistent with linkage mapping data from *Lactuca* that indicated the Compositae WGD was actually a triplication (Truco et al., 2013). The two rounds of WGD observed in our MAPS analyses is consistent with a shared paleotetraploidization in the MRCA of the Compositae and Calyceraceae with a nested paleohexaploidization in the ancestry of the core Compositae.

## DISCUSSION

Our analyses reveal a complex history of paleopolyploidy in the largest family of flowering plants. Single species (Ks plots) and multi-species phylogenomic analyses (MAPS) both place the previously identified Compositae (Asteraceae) WGD (Barker et al., 2008) in the ancestry of the Compositae and its sister family Calyceraceae. We did not find evidence of the WGD in any of the more closely related families of flowering plants, such as Goodeniaceae, Menyanthaceae, and Campanulaceae. However, we uncovered evidence for a second round of polyploidy among the core Compositae we analyzed, including representatives of *Helianthus, Lactuca, Artemisia,* and *Centaurea.* Although not readily apparent in our Ks plots, our phylogenomic approach is able to disentangle the different duplication events in the nuclear gene family phylogenies. Notably, our Ks plots and MAPS analyses did not find evidence of a more recent WGD in *Artemisia* (Anthemideae) and support our previous placement of the putative Heliantheae WGD (Baldwin et al., 2002; Barker et al., 2008). Given the phylogenetic position of these inferred WGDs, our results indicate that most species of Compositae experienced at least two rounds of paleopolyploidy in the last ~50 million years.

The phylogenetically nested WGDs are consistent with our expectations for a paleohexaploidy in the ancestry of the Compositae. A high density linkage map of *Lactuca sativa* had previously found that the lettuce genome was triplicated (Truco et al., 2013). In nature, hexaploids are not formed from a single, instant triplication event, but rather consecutive genome duplications (Harlan and DeWet, 1975; Ramsey and Schemske, 1998). For a paleohexaploidy, the consecutive duplications appear as triplications in linkage map or syntenic data. However, it is difficult to distinguish these independent events in Ks plots because the duplications often occur too close in time to resolve, especially after many millions of years of evolution. If extant lineages survive from both duplication events, then it may be possible to resolve the two rounds of duplication in gene family phylogenies. That appears to be the case in the Compositae. Our analyses indicate that the Calyceraceae and at least tribe Barnadesieae descend from the first round of WGD, whereas the more derived Compositae in our analyses have a second polyploidy in their ancestry. Although we cannot resolve the exact mechanism of hexaploid formation with our present analyses, we can assign putative ploidal levels to these WGDs. The total evidence available from our present analyses and previous research (Barker et al., 2008; Truco et al., 2013; Scaglione et al., 2016)indicates that the Calyceraceae and Barnadesieae are likely descendants of a paleotetraploid and the more derived Compositae tribes have a paleohexaploid ancestor. Previous analyses of the tribe Brassiceae (Lagercrantz and Lydiate, 1996; Lysak et al., 2005; Tang et al., 2012), the Solanaceae (Tomato Genome Consortium, 2012), and the eudicot triplication (Tang et al., 2008; Jiao et al., 2012; Vekemans et al., 2012) have used various gene family approaches to characterize these paleohexaploidies. There is no evidence in these cases for extant descendants of the paleotetraploid lineage. The Compositae and Calyceraceae represent a unique biological example with surviving lineages of both the paleotetraploid and paleohexaploid ancestors.

Although not precisely placed in the Compositae phylogeny because of limited sampling, we find that the second round of WGD may co-occur with many significant events in the evolution of the Compositae. The paleohexaploidization occurs in the same region of the phylogeny as two chloroplast DNA inversions (Jansen and Palmer, 1987; Kim et al., 2005). These inversions arose simultaneously, or nearly so, and are found among all sampled Compositae except *Barnadesia.* Intriguingly, our second WGD is co-located in the same region as these classic chloroplast inversions. It may be that the plastid inversions were introduced from a related species during an ancient hybridization event that produced the paleohexaploid ancestor. Two different plastid ancestors of extant Compositae may explain the rapid and simultaneous appearance of these plastid inversions in the phylogeny. Ancient hybridization and hexaploidization may have captured a plastid lineage that would otherwise have gone extinct. At present, we do not have sufficient resolution from our nuclear transcriptomes to assess if the paleohexaploid was an allo‐ or autohexaploid, but such diagnoses will be an interesting area of future research. Plastid inversions and other rearrangements have been widely used to define major lineages of plants (Downie and Palmer, 1992; Doyle et al., 1992; Raubeson and Jansen, 1992; Graham and Olmstead, 2000; Schwarz et al., 2015). It is possible that ancient allopolyploidy in the Compositae and other clades may have captured these chloroplast genome variants. Combining our nuclear genome based placements of paleopolyploidy with abrupt chloroplast rearrangements as well as novel mitochondrial gene transfers (Adams et al., 2000, 2001, 2002; Adams and Palmer, 2003; Brandvain et al., 2007) may be a useful approach to recognize potential paleoallopolyploidies.

The inferred paleohexaploidy also co-occurs with the evolution of Compositae tribes outside of South America. Evidence suggests that the Calyceraceae and Composite likely originated and initially diversified in present-day South America (Panero and Funk, 2008; Funk et al., 2009; Barreda et al., 2010). The early branching lineages of Compositae, such as Barnadesieae and Mutisieae, are largely restricted to South America. Most of the family’s diversity is found among the tribes that left South America and evolved on other continents (Panero and Funk, 2008). All of these “out of South America” tribes, which account for 96% of the family’s diversity, are descended from the inferred paleohexaploid ancestor. In contrast, most of the South American lineages are descended from the paleotetraploid ancestor. It is unclear from our present analyses whether the genetics of the paleohexaploidy contributed to increased net diversification, facilitated intercontinental movement and ecological expansion, or if the association is simply a coincidence in a large phylogeny. Further sampling will permit more precise placement of these WGDs to better evaluate these polyploidies and diversification of the Compositae. The survival of paleotetraploid and paleohexaploid lineages represents a unique opportunity to study the evolutionary outcomes and consequences of polyploidy of different ploidal levels.

An immediate observation from our results is that the Compositae WGD is not perfectly correlated with the origin or diversification of the Compositae. A recent analysis found that our previously identified Compositae WGD (Barker et al., 2008) was “perfectly associated” with an increase in net diversification rate (Tank et al., 2015). Given the information available at the time and the density of sampling in Tank et al. (2015), this is not an inaccurate statement. However, our new results suggest that the initial paleotetraploidy may have little association with increased net diversification as the Calyceraceae consists of only ~25 species and only ~4% of Compositae species are among the early diverging South American lineages (Panero and Funk, 2008; Funk et al., 2009). The inferred paleohexaploidy is correlated with a previously identified increase in net diversification rate that occurred just after the divergence of Barnadesioideae from the more derived Compositae (Smith et al., 2011). This location in the Compositae phylogeny is a “hot spot” of activity: plastid inversions, global expansion of the family, and now a paleohexaploidy. Determining if and how these events are related and ultimately influenced the diversity of the Compositae will require more transcriptomic and genomic sampling of the early diverging lineages. Although our analyses do not support a “lag” between the paleohexaploidy and diversification of the Compositae (Schranz et al., 2012), the nature of the relationship between polyploidy and diversification of the largest family of flowering plants remains unclear. Analyses of diverse Compositae genomes will permit us to test if increased divergent resolution of genes (Werth and Windham, 1991; Lynch et al., 2000; Scannell et al., 2006; McGrath et al., 2014; Muir and Hahn, 2015), co-evolutionary innovations driven by duplication (Edger et al., 2015), or other mechanisms of post-polyploid genome evolution (Arrigo and Barker, 2012; Garsmeur et al., 2014) contributed to the diversification of the Compositae. Ultimately, functional analyses connecting WGD duplicated genes with key morphological traits of the Compositae, such as Chapman et al. (2008), are needed to reveal the contribution of paleopolyploidy to plant diversity. Considering that as much as 40% of the genes in species of Compositae are derived from these WGDs (Barker et al., 2008), genomic analyses may provide new insight into how genetic diversity generated by paleopolyploidy has contributed to the domestication and diversity of crops (Hodgins, Lai, et al., 2014; Renny-Byfield and Wendel, 2014) as well as the evolution of weedy species in the family (Lai, Kane, et al., 2012; Dlugosch et al., 2013, 2015; Hodgins et al., 2013; Hodgins, Bock, et al., 2014). The complex WGDs in the Compositae and Calyceraceae provide a unique opportunity to explore these hypotheses.

---

Figure 1. Histograms of gene age distributions for six species of Compositae and related families. Y-axis is number of gene duplications; x-axis is synonymous divergence (Ks) of the duplication. Peaks of gene duplication in A, B, and C correspond to WGDs. A. *Artemisia annua* (Compositae); B. *Barnadesia spinosa* (Compositae); C. *Acicarpha spathulata* (Calyceraceae); D. *Scaevola aemula* (Goodeniaceae); E. *Nymphoides peltata* (Menyanthaceae); F. *Platycodon grandiflorus* (Campanulaceae). Image of *Sarnadesia* provided by Quentin Cronk. Platycodon image from www.biolib.de and licensed for re-use under the Creative Commons Attribution-Share Alike License. Images A, C, D, and E from Wikipedia and licensed for re-use under the Creative Commons Attribution-Share Alike License. *Artemisia* image by Kristian Peters, *Acicarpha* image by Marcia Stefani, *Scaevola* image by Nemracc, and *Nymphoides* image by TeunSpaans.

Figure 2. Species tree of Compositae and related families with gene duplications at each node as inferred by MAPS from across 1814 nuclear gene family phylogenies. The percentage of gene subtrees consistent with the species tree that support a shared gene duplication is indicated at each node. Numbers above nodes indicate number of subtrees that support a shared duplication out of the total number of subtrees consistent with the species tree. Colored circles highlight inferred WGDs at nodes N3 and N5. An inferred paleotetraploidy is inferred in the ancestry of Compositae and Calyceraceae at node N5. A second round of WGD is inferred at node N3 and corresponds to a putative paleohexaploidy in the ancestry of most Compositae.

Figure 3. A histogram of the percentage of gene duplications shared across 1,814 nuclear gene families inferred by MAPS at each node in our analyzed species tree. The node numbers correspond to nodes in Figure 2. Two WGDs are inferred at the two peaks of shared gene duplication at nodes N3 and N5.

